# Interaural time and level differences contribute differently to source segregation and spatial selection

**DOI:** 10.1101/2025.10.30.685652

**Authors:** Benjamin N. Richardson, Sahil Luthra, Jana M. Kainerstorfer, Barbara Shinn-Cunningham, Christopher A. Brown

## Abstract

Interaural time and level differences are crucial in sound localization, yet their contributions to sound source segregation and spatial selection remain underspecified. Here, participants completed a spatial auditory selective attention task while we measured hemodynamic activity in the prefrontal cortex and superior temporal gyrus using functional near-infrared spectroscopy. Participants listened to a target sound stream and a simultaneous spatially separated speech stream or white noise masker. Sound streams were spatialized with either 50 μs interaural time differences (ITDs), 500 μs ITDs, naturally occurring interaural level differences (ILDs) from a non-individualized head-related transfer function (HRTF), or broadband 10 dB ILDs. Behavioral results revealed a stronger effect of spatial cues when the masker was speech. Error patterns differed in the two difficult conditions, small ITDs and natural ILDs: Small ITDs produced lower hit rates, while naturally occurring ILDs produced higher false alarm rates. Small ITDs led to greater activity in prefrontal cortex and activity in superior temporal gyrus that was lateralized, greater in the hemisphere contralateral to attentional focus, consistent with previous reports. These results suggest that natural ILDs alone support source segregation even if they are insufficient to cause large shifts in perceived lateralization, explaining high false alarm rates (confusions between target and distractor words). In contrast, small ITDs alone may be insufficient to segregate competing sources, leading to low hit and false alarm rates. Together, these results reveal differences in how ITDs and ILDs contribute to auditory scene analysis and spatial attention.

## Introduction

In many typical sound environments, the signals reaching a listener’s ears comprise a rich mixture of concurrent, spectro-temporally overlapping but independent sounds, often arriving from different directions (Cherry, 1953). To understand a sound in such settings, the brain must achieve at least two interrelated tasks: 1) *segregate* the sources in the mixture, forming distinct perceptual objects by determining what acoustic energy belongs to what source and 2) *select* which object to process in detail by focusing selective attention to filter out competing sources (Noyce et al., 2023; Shinn-Cunningham, 2008a). Many acoustic factors influence both segregation and selection, including pitch, temporal envelope cues, and spatial features, among others (Middlebrooks and Waters, 2020; Moore and Gockel, 2002). The current work examines how spatial cues influence segregation and spatial selection.

Spatial cues are relatively weak compared to spectrotemporal cues in grouping together simultaneous sound elements, but they play a larger role in “streaming” sounds across silent gaps (Ihlefeld and Shinn-Cunningham, 2008; Middlebrooks, 2017; Moore and Gockel, 2002). Perceived spatial separation can also guide spatial attention to a target in a mixture (Arbogast et al., 2002; Freyman et al., 1999; Kidd et al., 2010). Yet spatial cues have little influence if competing sounds have different and distinct spectrotemporal structure, such as when a woman speaks over a background of radio static. When spectrotemporal features alone are enough to support segregation and selection, spatial separation between competing sources has little effect on psychophysical outcomes (Culling and Summerfield, 1995; Darwin, 2007; Kubovy, 1988; Kubovy and Van Valkenburg, 2001; Noyce et al., 2023). Thus, the impact of spatial separation on auditory attention depends critically on the spectrotemporal properties of competing sounds.

The influence of spatial cues on segregation and selection helps explain spatial release from masking (SRM), wherein listeners are better at detecting and understanding a target in the presence of a spatially separated masker compared to when they are co-located (Freyman et al., 1999; Middlebrooks and Waters, 2020). The most important and robust spatial cues for SRM are interaural time and level differences (ITDs and ILDs). These cues help listeners to selectively attend to a target, though their specific contributions to segregation versus selection remain underspecified.

ITDs and ILDs have distinct effects on the signals reaching the ears, which may impact how they contribute to SRM. Acoustically, ITDs do not alter the target-to-masker ratio (TMR) in the mixture of signals reaching the ears; thus, any benefits from ITDs must arise from explicit spatial computations within the auditory pathway. In contrast, ILDs affect the TMR in the mixture of signals reaching each ear; typically, when a target and a masker have different ILDs, the TMR will be larger at the ear nearer the target, producing a “better ear” effect (Bronkhorst and Plomp, 1988; Glyde et al., 2013a). Perceptually, this TMR boost can improve target understanding even in the absence of perceived spatial separation or in monaural listening conditions by simply reducing energetic masking. Physiologically, a source containing ILDs is represented asymmetrically in auditory cortex including the superior temporal gyrus or STG), with a stronger representation in the hemisphere contralateral to the ear receiving the more intense signal (Higgins et al., 2017). A direct consequence of this cortical lateralization is that the neural representations of competing sounds with opposing ILDs show inherently less overlap than representations of sources containing only ITDs. This lateralization may support perceptual source segregation even in the absence of explicit spatial effects. These differences suggest that ITDs and ILDs may support SRM through different mechanisms.

The extent to which ILDs support SRM will also depend on the spectrotemporal properties of competing sounds. Natural ILDs are substantial only in high frequencies (Yost and Dye, 1988). Because high-frequency components of speech group with their corresponding low-frequency components (Best et al., 2007; Bregman, 1990; Darwin, 1997), natural ILDs may nevertheless help with perceptual segregation of competing speech. Yet when paired with uninformative or zero ITDs, the perceived location of speech containing only natural ILDs and no usable ITDs will be nearer midline, since ITDs at low and mid frequencies dominate lateralization (Best et al., 2007; Ellinger et al., 2017; Glyde et al., 2013b; Wightman and Kistler, 1992). As a result, competing sources with only natural, opposing ILDs may be less likely to be perceived as coming from distinct locations, making source selection difficult. In contrast, broadband ILDs—applied uniformly across frequencies—can provide strong grouping and separation cues.

Magnifying ILDs to larger-than-natural magnitudes can enhance SRM in both normal-hearing listeners and cochlear implant (CI) users (Brown, 2014; Richardson et al., 2025). Without ILD magnification, CI users typically do not show significant SRM, likely because cochlear implants do not preserve ITDs and compress ILDs (Brown, 2018; Dorman et al., 2014; Gray et al., 2021). The current study seeks to clarify how ILDs contribute to SRM in normal hearing listeners in order to begin to identify how enhanced ILDs benefit CI users.

Functional neuroimaging studies reveal that spatial attention engages a visuo-spatial control network that includes prefrontal cortex (PFC; (Kong et al., 2014; Michalka et al., 2015; Noyce et al., 2022) and enhances contralateral bias in the representation of sound in auditory cortex (Alho et al., 1998, 1999, 2003; Ciaramitaro et al., 2007; Jäncke and Shah, 2002; Yang and Mayer, 2014). Furthermore, previous functional near-infrared spectroscopy (fNIRS) studies show that PFC activity varies with cognitive demands, reflecting individual differences in speech intelligibility (Lawrence et al., 2018; Zhou et al., 2022a, 2022b), perceived task difficulty (Zhou et al., 2022a, 2022b), and spatial separation between target and masker sounds (Zhang et al., 2021a). Measures of effort, including hemodynamic activity in PFC, show a non-monotic relationship with task difficulty: Activity is larger when auditory attention is actively deployed compared to passive listening (Zhang et al., 2018), and reduced responses can reflect either very low or very high difficulty (Lawrence et al., 2018). In a similar spatial selective attention task to that used in the present study, PFC showed less activity when target and masker were dichotic than when lateralized with relatively small ITDs (Zhang et al., 2021a). This result could reflect reduced demands on spatial attention when target and masker locations are more perceptually distinct (infinite ILDs compared to small ITDs); however, it could also reflect reduced effort expended to segregate sources lateralized with ILDs than with ITDs. Specifically, because neural representations are automatically lateralized for sources with ILDs, computational demands of segregation and selection may be reduced, resulting in decreased activation in PFC.

Previous investigations form a general picture of how ITDs and ILDs contribute to segregation and spatial selection. ITDs tend to dominate lateralization, and ILDs as they naturally occur are small at frequencies containing speech energy. ILDs do, however, result in the preferential representation of a sound in the contralateral auditory cortex. This separation of neural populations that represent competing sounds, along with top-down modulation from PFC, is required for successful selective attention. This has been explored using fNIRS imaging in both PFC and STG, but often separating competing sounds diotically (Zhou et al., 2022b) or using only ITDs (Zhang et al., 2018, 2021a). Here, we address this gap by comparing behavioral responses and neural activity in PFC and STG with either ITDs or ILDs during a spatial selective attention task.

In the current study, competing sounds were separated with small ITDs, large ITDs, natural ILDs, or broadband ILDs under conditions where the masker was either competing speech or white noise. Participants responded to target color words in one stream. We had several hypotheses about how the masker type (speech, noise) and spatial cues would interact to affect segregation and spatial selection in behavioral performance. First, we expected that a speech masker would more readily mask the target speech sound than a noise masker. Because of a lack of segregation cues in this case, participants would more heavily rely on spatial cues (Culling and Summerfield, 1995; Darwin, 2007; Kubovy and Van Valkenburg, 2001; Noyce et al., 2023): Larger spatial separation would engender better performance as indexed by hit rate and false alarm rate. Second, we hypothesized that conditions that provided small spatial cues in low frequencies would result in greater listening difficulty, reflected in poorer performance. That is, we expected worse behavioral performance with small ITD cues compared to large ITD cues, and worse performance for natural ILDs compared to broadband ILDs. Of particular interest is how these performance improvements might manifest with respect to hit rates and false alarms. For instance, if a change in spatial cues were to induce an increase in hit rates but also in false alarms, we would infer that listeners are able to accurately segregate the two streams (leading to enhanced hit rates) but that they struggle with selection (leading to them falsely respond to color words in the non-target stream). Accordingly, we view hit rates and false alarm rates in this task as indices of segregation and selection, respectively.

We further hypothesized that greater listening effort, as reflected in performance, would also manifest in greater activity in PFC. Specifically, we expected that small ITDs would drive greater PFC engagement relative to large ITDs, and natural ILDs than broadband ILDs. Finally, we predicted that STG activity would lateralize contralateral to the attended direction, consistent with previous reports (Alho et al., 1999, 2003; Ciaramitaro et al., 2007; Jäncke and Shah, 2002; Yang and Mayer, 2014).

## Materials & Methods

### Participants

Thirty native English speaking listeners participated in this experiment (27 female, 3 male, age 22.6 ± 4.45 S.D.). Although our sample was predominantly female and sampled from university students, we did not anticipate that there are any gender differences in performance on this psychometric task. All statistical analyses were conducted within-subject. All participants had normal hearing, defined as having pure-tone audiometric thresholds of ≤ 20 dB HL at octave spaced frequencies from 250 Hz to 8 kHz. All listeners gave written informed consent prior to participating in the study. All testing was administered according to the guidelines of the Institutional Review Board of the University of Pittsburgh.

### Overview of the Task

Subjects completed a color-word detection task in a selective attention paradigm with two competing, spatially separated sound streams. Figure 1 shows a schematic of target and masker streams. The target stream consisted of temporally jittered words spoken by a single male talker. The competing stream was either speech from the same talker or white noise. Within an experimental block, target and masker sequences were spatialized such that they were perceived to originate from opposing hemifields, one on the left and the other on the right, assigned randomly on each block.

**Figure 1.**
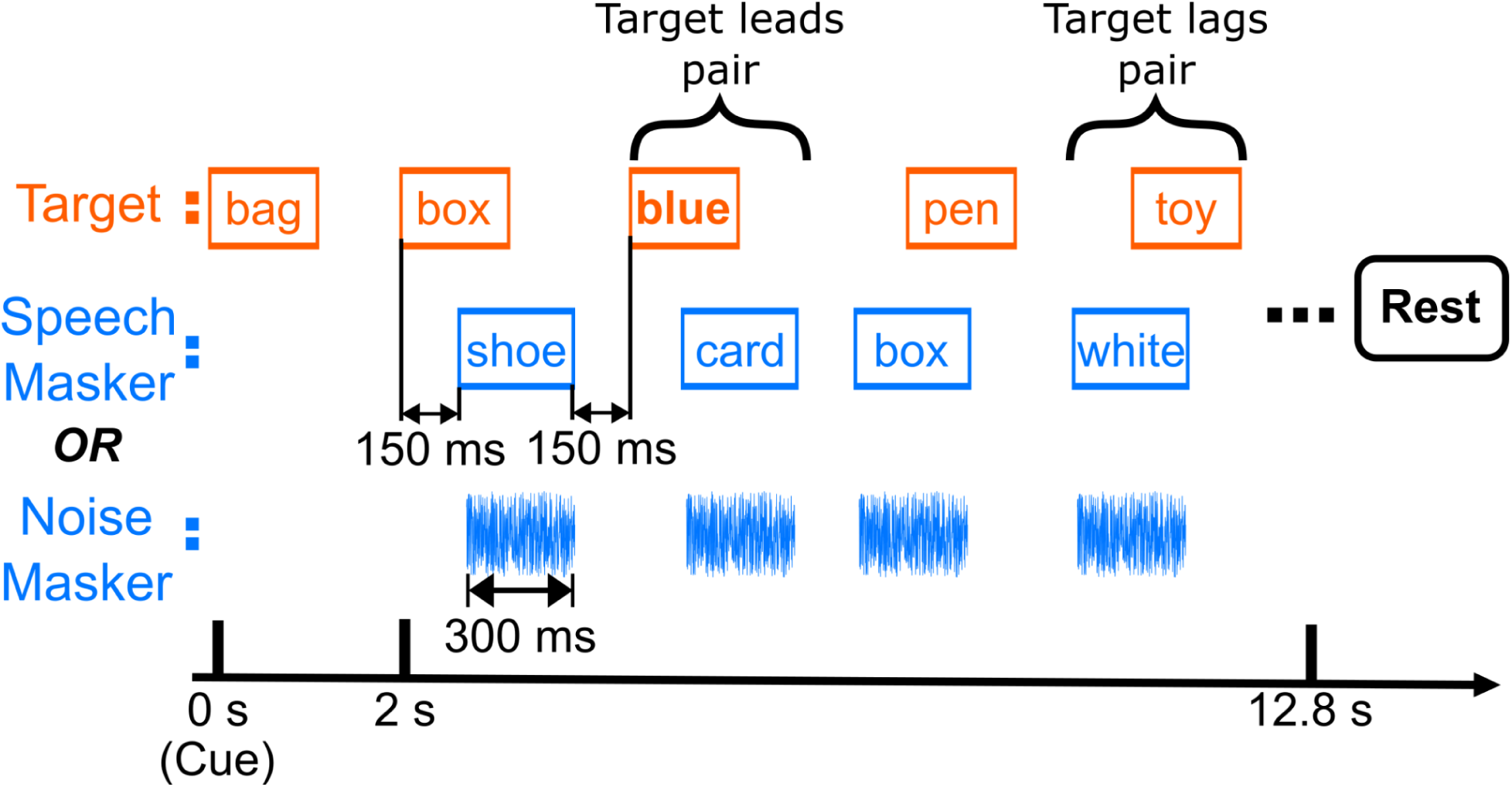
Schematic of a single block. Listeners first heard a cue word “bag” from the target stream. Then, they responded by button press to color words in the target stream (top row). The target stream was presented against either a speech or noise masker at the same intensity (i.e., with a 0-dB target-to-masker ratio). Each block was followed by 13-18 second rest.

For the speech masker condition, target and masker sequences each consisted of 18 word tokens, which were presented in pairs (one target token and one masker token, see *Stimuli*). A word token pair occurred every 600 ms. Within each token pair, the two 300-ms-long tokens partially overlapped in time, with the second one starting 150 ms after the first. To ensure that listeners relied on spatial attention rather than temporal regularity, which token in a pair occurred first was randomly selected, independently, for each pair in a block. A 150-ms-long silent gap followed the second token before the next pair began. For the noise masker condition, masker word tokens were replaced by white noise bursts.

Listeners were instructed to press a key on a handheld keypad whenever they detected a color word within the target stream. Data collection was divided into blocks, each of which presented one of the two different masker streams (speech or noise; see Fig. 1) and one of four different spatialization conditions, described in detail below (see Stimuli). At the start of each block, a single presentation of the word “bag” was presented from the side of the target stream to cue the participant as to which direction to attend. Two seconds after the start of the cue word, the target and masker streams began playing. Thus, each block consisted of the 2-sec cue interval, followed by 18 token pairs, each of which was 600 ms long, for a total block duration of 12.8 s (see Figure 1).

Each unique combination of masker type and spatialization condition was presented in 8 different blocks, yielding a total of 64 blocks (8 blocks for each of 2 masker types x 4 spatializations). In addition, for each masker and spatialization condition, 4 blocks presented the target stream to the left (masker to the right), and 4 had the target to the right (masker to the left). The order of these 64 blocks was randomized separately for each participant. A silent period followed each block, with the duration selected randomly from a uniform distribution from 13-18 seconds. This silent period allowed hemodynamic responses to return to baseline (see *fNIRS Data Acquisition and Analysis*). Subjects were given the option to pause after each block and were prompted to take a break at the halfway point of the experiment.

### Stimuli

All stimulus processing was completed in the Python programming language (Van Rossum and Drake, 2009), with signal processing functions that were either from the *scipy* package (Virtanen et al., 2020) or custom written.

Target tokens were American English words. These speech stimuli were constructed from a list of 16 words, recorded in isolation by a single male talker (Kidd et al., 2008). The words were selected from a set of 12 object words: <*hat*, *bag*, *card*, *chair*, *desk*, *glove*, *pen*, *shoe*, *sock*, *spoon*, *table*, *toy*> and a set of four color words: <*red*, *white*, *blue*, *green*>. All word tokens were edited to have a duration of 300 ms by using a waveform editor to isolate each token and the soundstretch synchronous-overlap-and-add algorithm to adjust duration (“SoundStretch Audio Processing Utility,” n.d.). Finally, tokens were equated in root-mean-square amplitude.

Target and masker word sequences were generated by randomly choosing 18 words, with replacement, from the object and color word sets. Each target stream included a minimum of 3 and a maximum of 5 color words; the other words were all from the object set. The number of color words in the masker stream was also drawn randomly between 3 and 5 words. The temporal position of color words was restricted such that color words could not occur within 2 indices of each other in either stream.

Two types of maskers were used: speech (second row, Figure 1) and noise (third row, Figure 1). Speech masker sequences were generated identically to target sequences, but with the additional constraint that within each target-masker pair, the masker word differed from the target word. Note that color words could occur in the speech masker sequence, but the listener was asked to ignore these “foil” color words. The speech masker contained words drawn from the same set (i.e., they were spoken by the same talker). This choice and the lead-lag jittered timing resulted in stimuli in which target and masker could only be differentiated based on their perceived directions: Participants did not have access to pitch differences, word-level timing cues for the task, or other features that could be used to determine whether a word was from the target or the masker.

Noise masker streams comprised independent 300-ms white noise bursts. The noise tokens were each ramped with 20 ms raised cosine onset and offset ramps. Each noise burst was flat spectrum broadband white noise and sampled independently, and so covered frequencies 0-22 kHz. Noise bursts were subjected to the same timing constraints as speech masker word tokens (Fig. 1). Color words across both target and speech masker streams were constrained to ensure that color words could not occur in successive word token pairs. The presentation level of each stream was set to 65 dB SPL resulting in a 0 dB target-to-masker ratio prior to applying spatial cues.

The four different spatialization conditions are illustrated in Figure 2. To each stream (target and masker, spatialized to opposite lateral locations), we either applied ± 50 μs ITD (“Small ITD”), ± 500 μs ITD (“Large ITD”), a naturally occurring ILD corresponding to 70 degrees azimuth (“Natural ILD”), or a 10 dB broadband ILD (“Broadband ILD”). To apply ITDs, the signal in the ear contralateral to the apparent source location was delayed in time by the appropriate number of samples (corresponding to 50 μs or 500 μs). In the natural ILD condition, the magnitude spectrum from a head-related transfer function (HRTF) for 70° azimuth, measured on a KEMAR manikin, was applied to spatialize sources to the right (Gardner and Martin, 1995). For sources to the left, the left and right magnitude spectra from this HRTF were swapped. The 70° location was chosen because it yielded the greatest average broadband ILD magnitude, equal to about 16 dB. In the broadband ILD condition, the signal in the contralateral ear was attenuated by 10 dB (equally across all frequencies). After spatialization, target and masker streams, one from the left and one from the right, were summed to create left and right channels.

**Figure 2.**
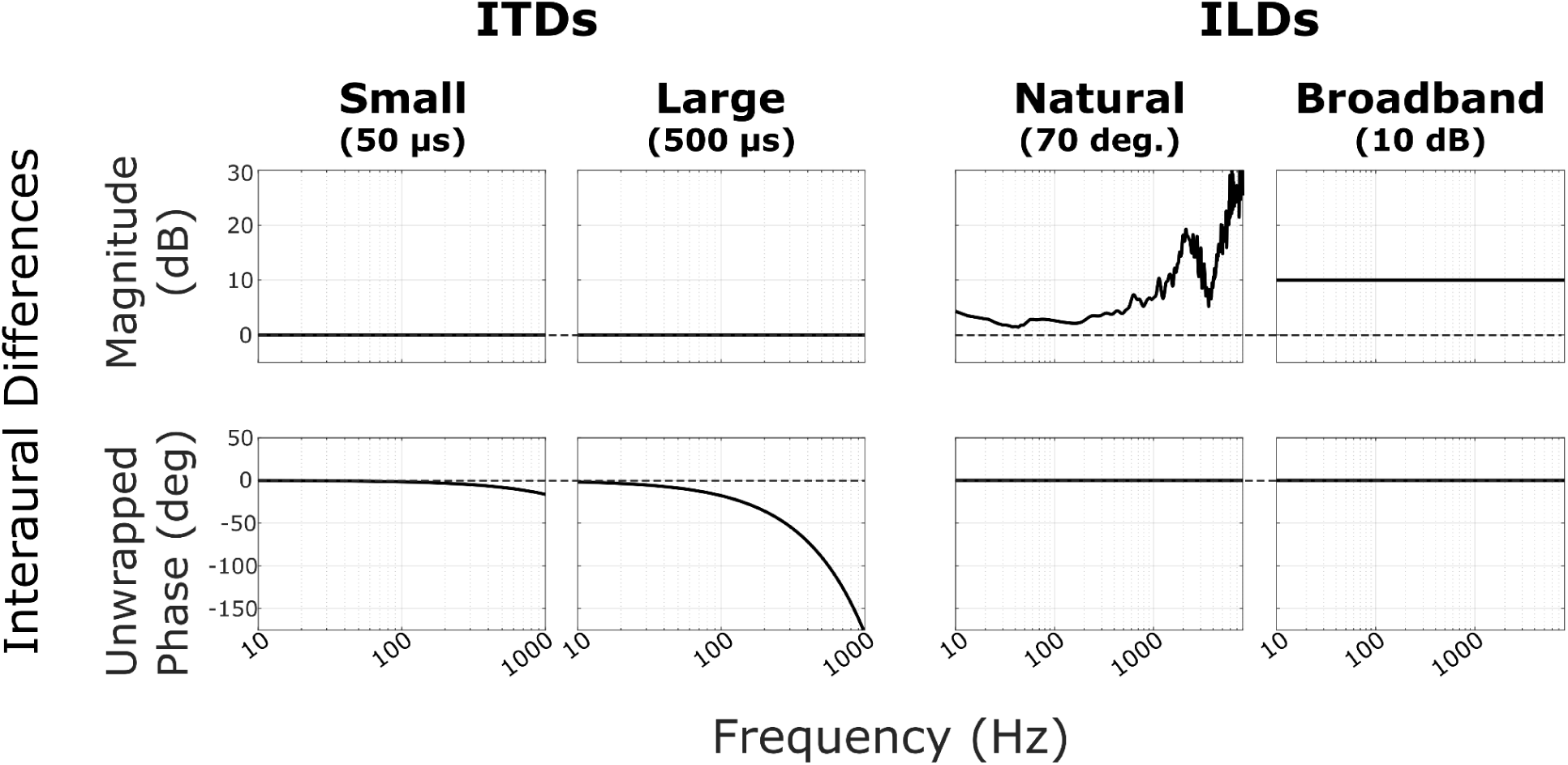
Spatialization conditions. The left (L) and right (R) ear signals for a given stream were spatialized with a 50 μs ITD (“Small ITD”), 500 μs ITD (“Large ITD”), naturally occurring ILD (HRTF at 70° azimuth, “Natural ILD”), or a 10 dB attenuation (“Broadband ILD”). Each subplot shows the magnitude (top row) and unwrapped phase (bottom row) properties of the interaural differences used. ITDs were implemented by pure time delay of the contralateral ear. Note that for the ITD plots, the unwrapped phase plots show a characteristic falloff with frequency proportional to 1/f, which correspond to group delays of 50 μs and 500 μs for small and large ITDs, respectively.

### Experimental Procedure

Participants were seated in an anechoic chamber, fitted with Etymotic ER-3a insert phones, and provided with a numeric keypad. Before the start of the experiment, subjects performed practice runs to familiarize themselves with the stimuli and task. Practice runs presented only a diotic target sequence with no masker and no spatial cues applied. Participants practiced pressing the keypad “enter” button when they heard a color word from the talker. Participants had the opportunity to discuss the task with the experimenter between practice runs. At least three practice runs were presented to each participant; however, each participant could perform as many practice runs as they needed to feel comfortable with the task. Participants were then instructed that there would be two streams during testing, each coming from a different location, and that the cue word ‘bag’ would be presented at the start of each block to indicate the location of the target sequence in that block.

### Behavioral Performance Estimation

In each block, we recorded the time of each button press and compared it to the onset of word tokens to determine rates of “hits” (correct responses to a target color word) and, for cases with the speech masker, “false alarms” (incorrect responses following a color word in the distractor stream). For each of these two response types, we defined windows for scoring response rates. “Hit windows” encompassed 300-1400 ms following the onset of each color word in the target stream; “false alarm windows” spanned 300-1400 ms after each color in the masker stream. To resolve ambiguity due to overlap of these potential windows, if a potential hit window overlapped a potential false alarm window, both the hit and false alarm windows were removed.

Additionally, if two potential time windows of the same type overlapped, the first of the two was kept and the subsequent window thrown out. Using this approach, all response windows were of equal length in time and did not overlap with one another. Since these windows were defined independent of when button presses occurred, they allowed us to operationally define hit rates and false alarm rates in a fair and unbiased manner. Finally, any button-press responses not associated with a color word window (whether or not that window was thrown out) were termed “random responses.”

We computed hit rates and random response counts for both masker types; we computed hit rates separately for lead and lag color word positions. Overall, few responses were removed due to overlapping hit and FA windows, although more were removed for a speech masker than a noise masker (mean 3.21 ± 0.20 S.D. clicks for speech masker; mean 0.00 ± 0.00 S.D. clicks for noise masker). Very few random responses (i.e., those that fell completely outside any color word response window, including deleted ones) occurred in any condition (mean 0.317 ± 0.053 S.D. clicks for speech masker; mean 0.283 ± 0.050 S.D. clicks for noise masker).

For the speech masker, we also calculated false alarm (FA) rates. We opted to analyze hit and FA rates because false alarms did not exist in the noise masker case, as there were no color words in the noise masker stream. In analyzing hit rates and false alarm rates separately, we also tap into two different aspects of task performance. Hit rate reflects the ability to segregate target from masker; a failure to segregate the competing streams will lead to low hit rates, because none of the words will be intelligible. In contrast, false alarm rates reflect failures to differentiate target and masker word locations. If target and masker streams are perceptually segregated and sufficiently distinct in their perceived locations, listeners should be able to focus attention on the target stream and ignore the masker stream (not responding to color words from the wrong direction).

Hit and false alarm response rates were calculated as the total sum of button presses in each of the corresponding remaining time windows divided by the total number of those windows. Rates were computed individually for each subject in each condition.

### fNIRS Data Acquisition and Analysis

Functional near-infrared spectroscopy (fNIRS) was used to record changes in concentration of oxygenated and deoxygenated hemoglobin concentration during the task. We used a NIRx NIRSport 2 (NIRx, Berlin) system and recorded at 760 and 850 nm wavelengths with a source-detector distance of 3 cm. The source-detector montage and sensitivity map are shown in Fig. 3. Subjects wore an fNIRS head-cap fitted with light sources and detectors covering the prefrontal cortex (PFC) and superior temporal gyrus (STG) bilaterally, resulting in 14 source-detector pairs. Source and detector locations were chosen by selecting “dorsolateral prefrontal cortex” and “superior temporal gyrus” in the fNIRS Optodes’ Location Decider (fOLD) software (Zimeo Morais et al., 2018). Short channel detectors were attached to each source in order to record systemic physiological signals from the scalp.

**Figure 3.**
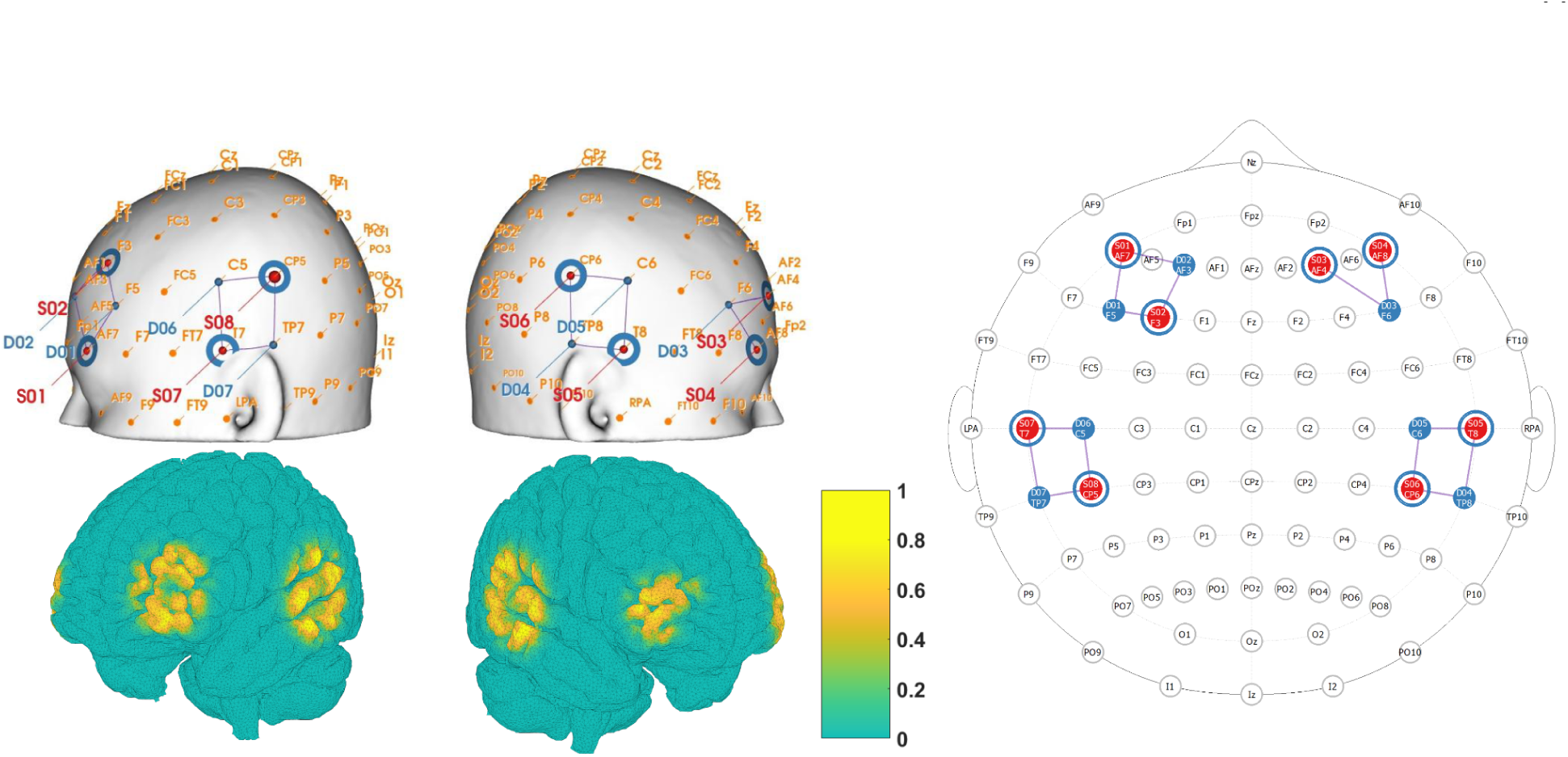
fNIRS montage and sensitivity map. Sensors covered the prefrontal cortex and superior temporal gyrus, bilaterally. Left: The top row shows placement of near-infrared light sources and detectors (end points of white lines), and the location of data channels (between each source and detector, yellow dots). The bottom row shows the normalized sensitivity map projected onto the ICBM 2009c nonlinear asymmetric brain model (Fonov et al., 2009). Right: 2D representation of source detector placement generated using the NIRSite montage creation software (NIRx).

fNIRS data processing was performed in Python using the MNE software package (Gramfort et al., 2013; Luke et al., 2021-4). Raw intensity values were first converted to optical density. To exclude source-detector pairs (channels) with poor data quality, we calculated the scalp coupling index (SCI) for each channel, a measure of the presence of the heart rate signal in the data (Pollonini et al., 2014). Channels with an SCI below 0.8 were removed from further analysis. Out of 14 long-distance channels, across the different participants an average of 1.42 ± 1.47 S.D. channels were removed due to low SCI. Next, high-frequency motion artifacts were removed using temporal derivative distribution repair (Fishburn et al., 2019). Time traces were then bandpass filtered between 0.01 and 0.3 Hz using a fourth order, zero-phase Butterworth filter to remove heartbeat and respiration signals. In order to further remove physiological artifacts, short channel information was regressed out of the optical density signal using the short_channel_regression function in MNE (Fabbri et al., 2004; Saager and Berger, 2005; Scholkmann et al., 2014). Finally, we applied the modified Beer-Lambert law (Kocsis et al., 2006) to the time traces of optical density to compute changes in oxygenated hemoglobin (ΔHbO) and deoxygenated hemoglobin (ΔHbR) concentrations.

The 13-18 second gap after each block ensured that hemoglobin levels returned to baseline before the next block began. The length of the silent period was chosen randomly from that range on each block in order to avoid the presence of a coherent signal in the fNIRS data at the rate of stimulus presentation. Raw fNIRS data for each block were epoched from −5 to 20 seconds from the onset of the sound cue. The first 5 seconds of each epoch (before and up to the cue) were used to set the baseline of ΔHbO and ΔHbR by subtracting the mean value over this period from the raw measure. Baselining was performed separately for ΔHbO and ΔHbR data. To quantify the strength of the hemodynamic response in each block, we calculated the mean ΔHbO value during sound stimulation (from stimulus onset to 12.8 seconds later) in each channel for each block.

### Statistical Analysis

We used linear mixed effects regression to analyze task performance and by-channel hemodynamic response magnitudes. All models were implemented in R version 4.2.1 (R Core Team, 2024) and fit using the “lmer” function in the lme4 library (Bates et al. 2015). To estimate main effects and interactions, we used the “mixed” function in the afex library (Singmann, H., Bolker, B., Westfall, J., & Aust, F., 2018), which uses likelihood ratio tests to assess the fit of a model with the effect of interest versus that of a simplified model without that effect. Thus, main effects and interactions are reported with chi-square values that index whether inclusion of the effect leads to a significant improvement in model fit. For all measures, we followed up on significant effects with Bonferroni-corrected pairwise t tests.

To analyze task performance, we fit separate models for hit rates and false alarm rates. For both analyses, we included fixed effects of spatialization (small ITD, large ITD, natural ILD, broadband ILD) and color word position (lead, lag), as well as random intercepts for each participant. We analyzed all spatialization conditions as levels of a single factor (rather than ITD/ILD as one factor and small/large as another) because ‘small’ and ‘large’ conditions within each cue type were not perceptually or physically equivalent; accordingly, potential interactions between cue type and cue magnitude would be misleading For the hit rate model only, we included a fixed effect of masker type (speech, noise). We did not include this effect for false alarms, since false alarms were not possible in the noise masker condition.

To compare hemodynamic response magnitudes, we fit separate statistical models for mean ΔHbO in PFC, mean ΔHbR in PFC, mean ΔHbO in STG, and mean ΔHbR in STG. We fit separate models for each masker type, given our a priori hypothesis that separate psychoacoustic mechanisms underlay performance with each masker type. For all fNIRS models, we included a fixed effect of spatialization, random intercepts for each subject, and random intercepts for each channel, allowing the model to account for baseline differences in activation across channels. Finally, for the STG models only, we also included a fixed effect of cortical hemisphere (contralateral, ipsilateral), driven by our a priori hypothesis that spatial selective attention will result in an increase in neural activity in the auditory cortex contralateral to attention (Alho et al., 2003; Ciaramitaro et al., 2007). We did not include a fixed effect of cortical hemisphere for the PFC data because the sensor montage was not symmetrical, and therefore did not image identical populations in left and right hemispheres.

For each behavioral and neural analysis, the omnibus model used sum-coded contrasts for all factors (e.g., the fixed effect of cortical hemisphere might be coded with a [1,-1] contrast for [contralateral, ipsilateral]). To perform pairwise comparisons between different levels of spatialization (e.g., small ITD vs. large ITD), we re-fit the omnibus model using treatment coding. With a treatment coding scheme, one level (e.g., small ITD) was set as a reference level, and beta estimates reflect pairwise comparisons between the reference level and each other level. To obtain all pairwise estimates, we ran four models, rotating which level was the reference level. Critically, these follow-up models do not differ in their fit to the data; all that differs across the models is the comparison captured by each beta coefficient. For all pairwise comparisons, we therefore report beta coefficients that capture the difference between those conditions in the same units as the dependent measure (e.g., hit rate).

## Results

### Hit rates varied between spatialization conditions

Hit rates across spatialization, masker type, and color word position are shown in the first column of Figure 4. The hit rates for the small ITD condition (leftmost data points) tended to be lower than for any other condition. Hit rates were slightly higher for the noise masker than the speech masker (compare bottom left and top left panels). Additionally, hit rates tended to be higher for a color word that occurred in the leading position within a word token pair than for a lagging position across all spatialization conditions (compare blue circles to corresponding yellow triangles).

**Figure 4.**
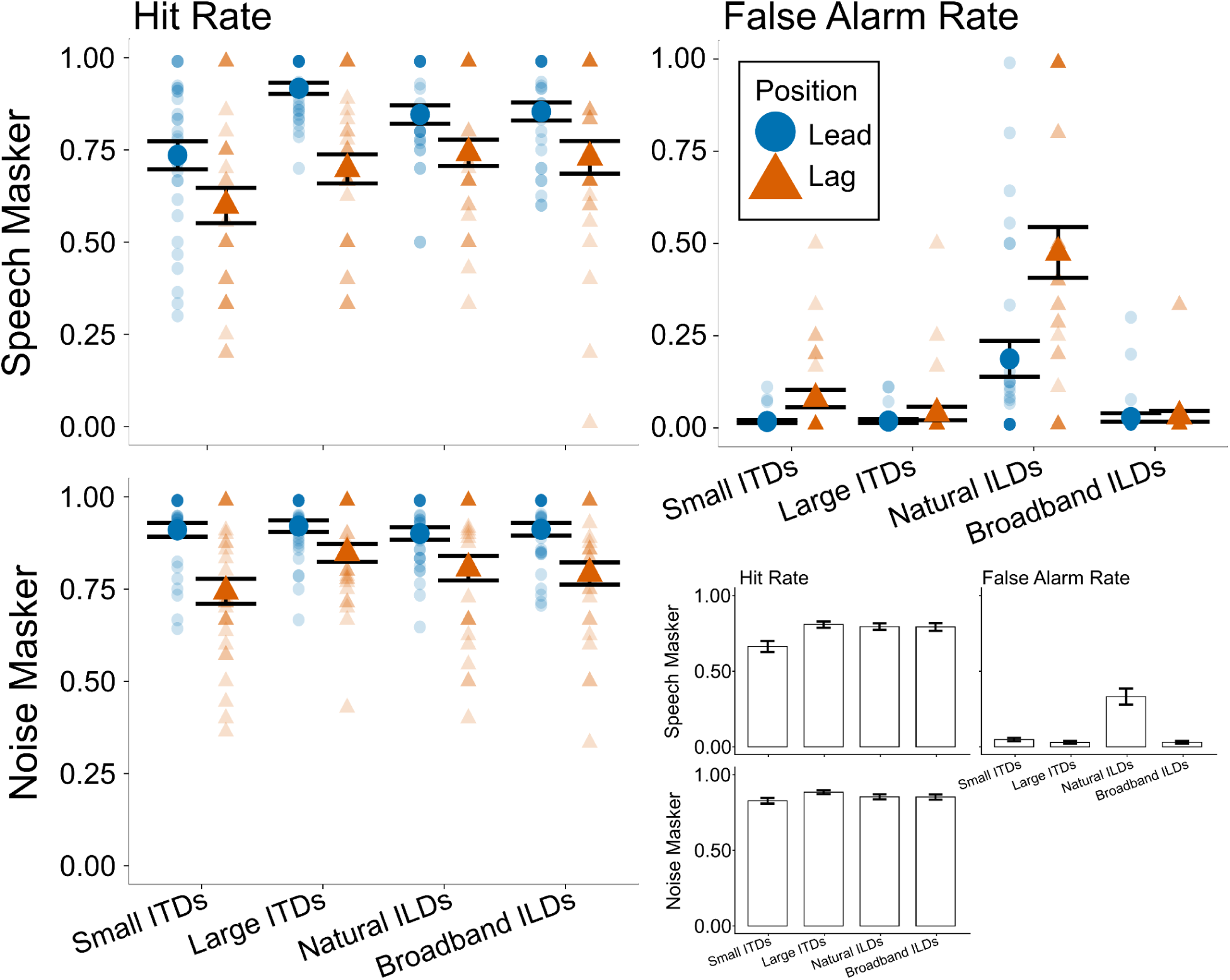
Hit and False Alarm Rates. First column (Hit Rates): The percentage of responses to color words in the target stream when the masker was speech (top row) or noise (bottom row). Second column, Top (False Alarm Rates): The percentage of responses to color words in the masker stream only when the masker was speech. For all plots, blue circles show results when a color word occurred in the leading position, and orange triangles show results for a color word in the lagging position. Small markers show individual participant data, and large markers show the mean across participants. Error bars indicate the mean and SEM across subjects in each condition. Second column bottom: The data represented in the other three subplots is collapsed across lead and lag word positions for ease of comparison. Bar height represents the mean proportion of responses in each category (hit rate, false alarm rate), and error bars show SEM across subjects in each condition.

Statistical analyses supported these observations. The linear mixed effects model revealed a significant main effect of masker type (χ^2^(1) = 37.17, p < 0.001), driven by higher hit rates with the noise masker (mean: 0.854) compared to with the speech masker (mean: 0.765). We also observed a significant main effect of color word position (χ^2^(1) = 75.26, p < 0.001), driven by higher hit rates for color words occurring in the leading position (mean: 0.875) compared to in the lagging position (mean: 0.745). The model also revealed a significant main effect of spatialization (χ^2^(3) = 26.40, p < 0.001) and a significant interaction between spatialization and masker type (χ^2^(3) = 8.35, p = 0.039). The three-way interaction between spatialization, masker, and color word position was not significant (χ^2^(3) = 5.45, p = 0.142).

To interpret the significant interaction between spatialization and masker type, we conducted post hoc separate analyses of the effect of spatialization within each masker type. Within the speech masker data, the hit rates with small ITDs (mean: 0.667) were significantly lower than with large ITDs (mean: 0.808; β = 0.140, p = 0.0001), natural ILDs (mean: 0.794; β = 0.127, p = 0.0005), and broadband ILDs (mean: 0.792; β = 0.125, p = 0.0006), but the large ITD, natural ILD, and broadband ILD hit rates did not differ significantly from one another. For the noise masker, the hit rates with small ITDs (mean: 0.827) were significantly lower than with large ITDs (mean: 0.884; β = 0.057, p= 0.034). Hit rates for all other pairs of conditions were not significantly different when the masker was noise.

### False alarm rates were larger with natural ILDs

As noted above, false alarm rates (reflecting responses to a color word in the distracting stream) could only be computed for the speech masker conditions (top row of Figure 4). As seen in the second column of Figure 4, the false alarm rates were near zero for small ITD, large ITD, and broadband ILD conditions; however, they were higher in the natural ILD condition, especially for lagging color words.

Statistical analyses confirmed these observations: The mixed effects model showed a significant main effect of spatialization (χ^2^(3) = 105.78, p < 0.001), a significant main effect of color word position (χ^2^(1) = 17.45, p < 0.001), and a significant interaction between the two (χ^2^(3) = 25.75, p < 0.001). To interpret the interaction, we conducted post hoc analyses across spatialization condition within each color word position. For the leading color word, false alarm rates were significantly larger in the natural ILD condition (mean: 0.187) than in the small ITD (mean: 0.018; β = −0.170, p < 0.0001), large ITD (mean: 0.019; β = −0.169, p < 0.0001), and broadband ILD (mean: 0.028; β = −0.159, p < 0.0001) conditions. An identical, but statistically stronger pattern arose for the lagging color word: False alarm rates were significantly larger in the natural ILD condition (mean: 0.476) than in the small ITD (mean: 0.079; β = −0.396, p < 0.0001), large ITD (mean: 0.040; β = −0.436, p < 0.0001), and broadband ILD (mean: 0.032; β = −0.444, p < 0.0001) conditions. None of the false alarm rates for leading and lagging color words in the small ITD, large ITD, or broadband ILD conditions differed significantly from one another.

### fNIRS Results

Figure 5 shows the block averaged ΔHbO (red lines) and ΔHbR (blue lines) mean time traces and standard errors across subjects (colored ribbons), averaged within each ROI and masker type. The left plot in each panel compares the small (solid) and large (dashed) ITD spatialization conditions, and the right plot compares natural (solid) and broadband (dashed) ILD spatializations. Hemodynamic responses in all regions and conditions show a canonical shape, with ΔHbO increasing with respect to baseline during sound stimulation (between 0 and 12.8 seconds) and decreasing thereafter (Luke et al., 2021-4; Quaresima and Ferrari, 2019).

**Figure 5.**
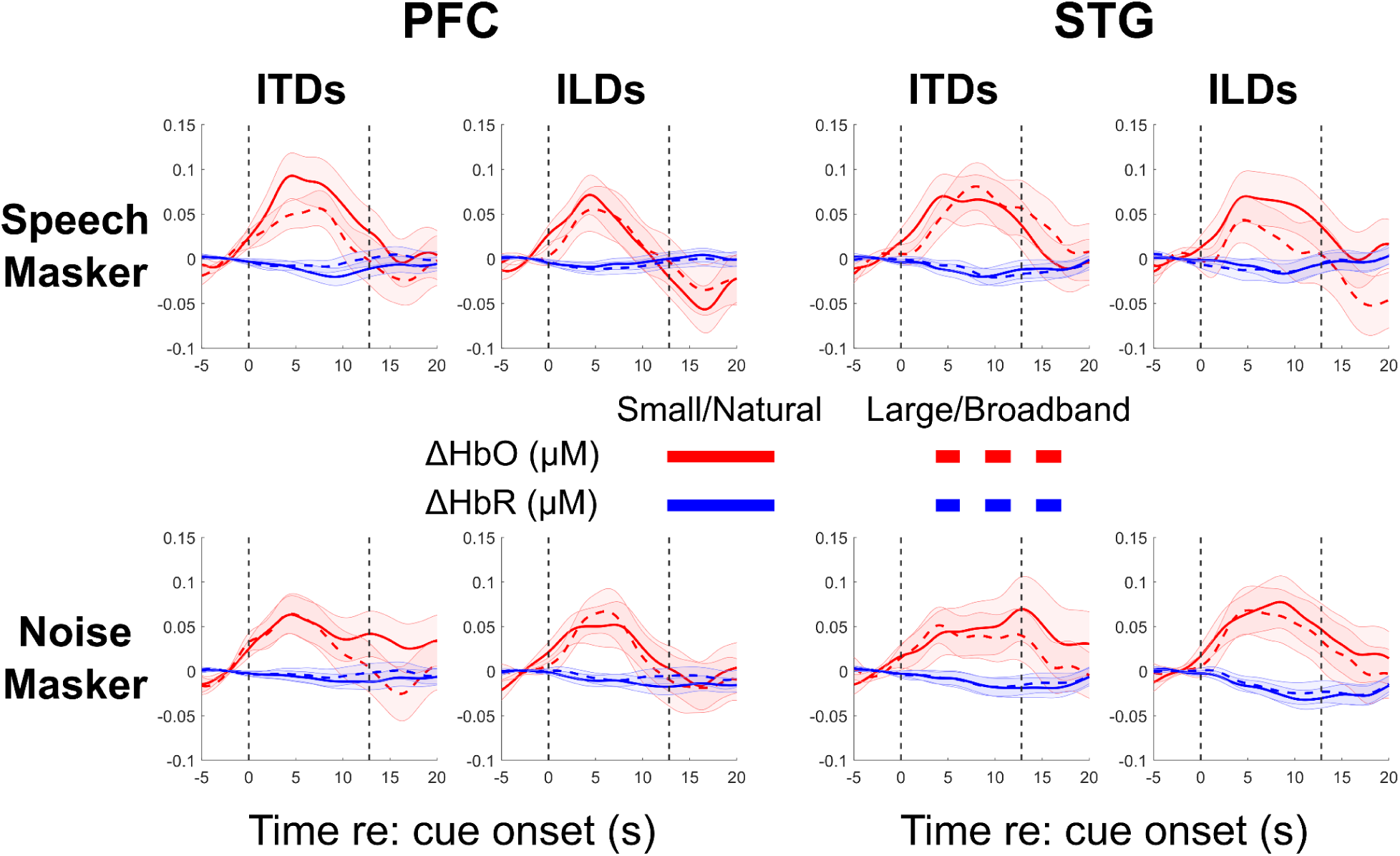
Block average hemodynamic responses. Each individual subplot shows the ΔHbO (red) and ΔHbR (blue) curves for each masker type and in each region. Shaded error bars show ± 1 S.E.M. Top left panel: Hemodynamic responses in PFC when the masker was speech (small vs. large ITD, left subpanel; Natural vs. Broadband ILD, right subpanel). Top right panel: as before, in STG when the masker was speech. Bottom left panel: as before, in PFC when the masker was noise. Bottom right panel: as before, in STG when the masker was noise.

Figure 6 summarizes the differences in hemodynamic responses by plotting the mean ΔHbO and ΔHbR averaged within prefrontal cortex during the block (0-12.8 seconds from cue onset). Mean ΔHbO for the small ITD condition was larger than for any other condition when the masker was speech, but mean ΔHbO did not vary strongly with spatialization when the masker was noise.

**Figure 6.**
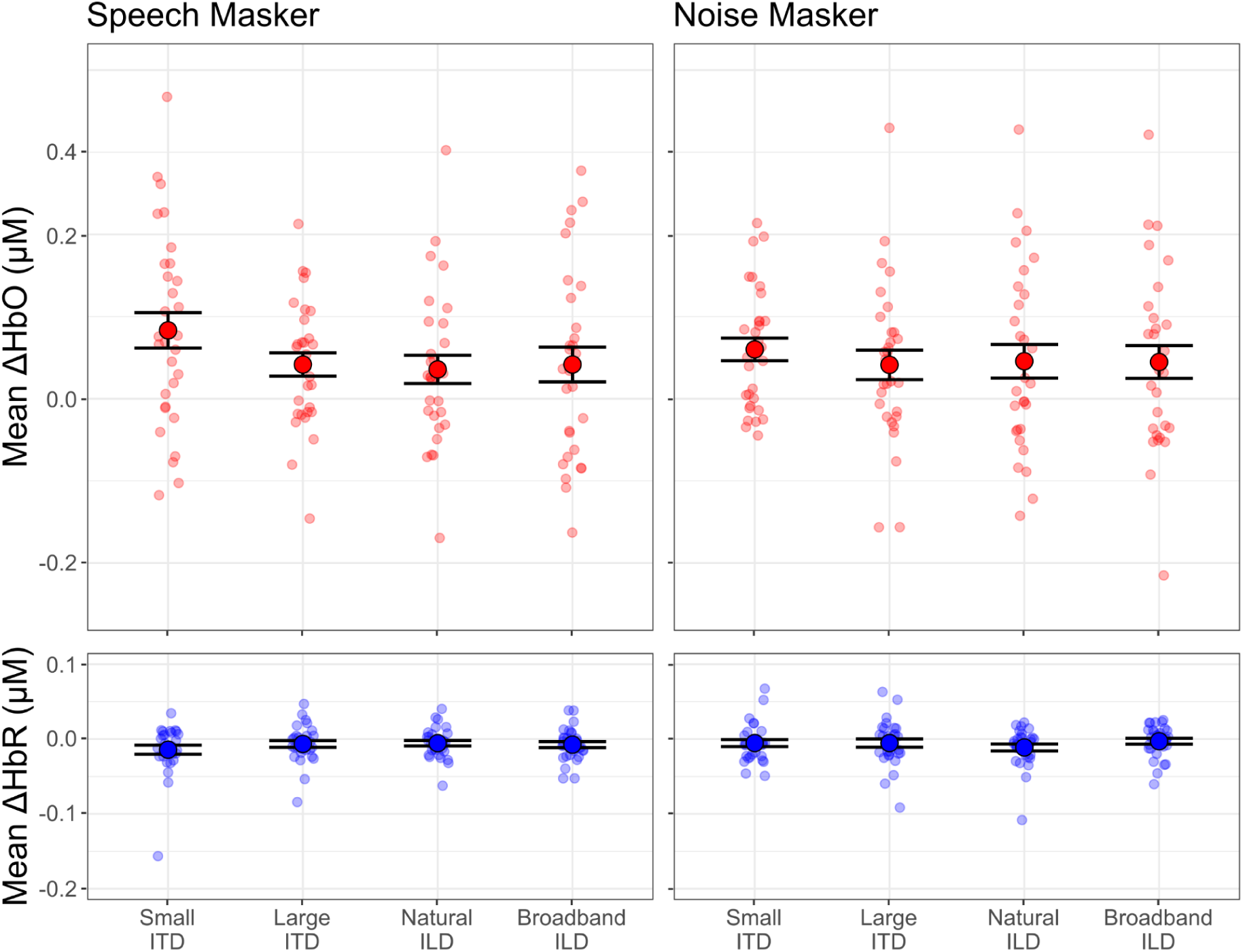
Mean hemodynamic response magnitudes in prefrontal cortex. Top row: Mean changes in oxygenated hemoglobin concentration (ΔHbO) when the masker was speech (left panel) and noise (right panel). Bottom row: Mean changes in deoxygenated hemoglobin concentration (ΔHbR) when the masker was speech (left panel) and noise (right panel). Small markers show individual participant data, and large markers show the mean across participants. Error bars indicate the mean and SEM across subjects in each condition.

Consistent with this summary, our statistical models only found a significant effect of spatialization in mean ΔHbO in PFC (χ^2^(3) = 27.15, p < 0.001). Post hoc tests showed this main effect was driven by significantly larger mean ΔHbO in the small ITD condition (mean: 0.086 μM) compared to the large ITD (mean: 0.039 μM; β = −0.047, p < 0.001), natural ILD (mean: 0.037 μM; β = −0.049, p < 0.001), and broadband ILD (mean: 0.044 μM; β = −0.042, p < 0.001) conditions, with no significant differences between the other three spatialization conditions. When the masker was noise, the effect of spatialization on mean ΔHbO in PFC was not significant (χ^2^(3) = 3.82, p = 0.282). Mean ΔHbR in PFC did not significantly vary in either the speech masker case (χ^2^(3) = 4.94, p = 0.176) or the noise masker case (χ^2^(3) = 4.48, p = 0.214).

Figure 7 shows the across-channel mean ΔHbO and mean ΔHbR in STG for each hemisphere (contralateral, ipsilateral to the attended direction). For the speech masker, hemodynamic responses in STG were similar across hemispheres and across spatialization cues for the large ITD, natural ILD, and broadband ILD conditions. However, in the small ITD condition, the hemodynamic response was stronger in the cortical hemisphere contralateral to the direction of attention. For the noise masker, hemodynamic responses varied more with spatial cue condition, but did not differ between hemispheres for large ITD or natural ILD conditions; however, for both small ITD and broadband ILD conditions, hemodynamic responses were larger in the hemisphere contralateral to the direction of attention.

**Figure 7.**
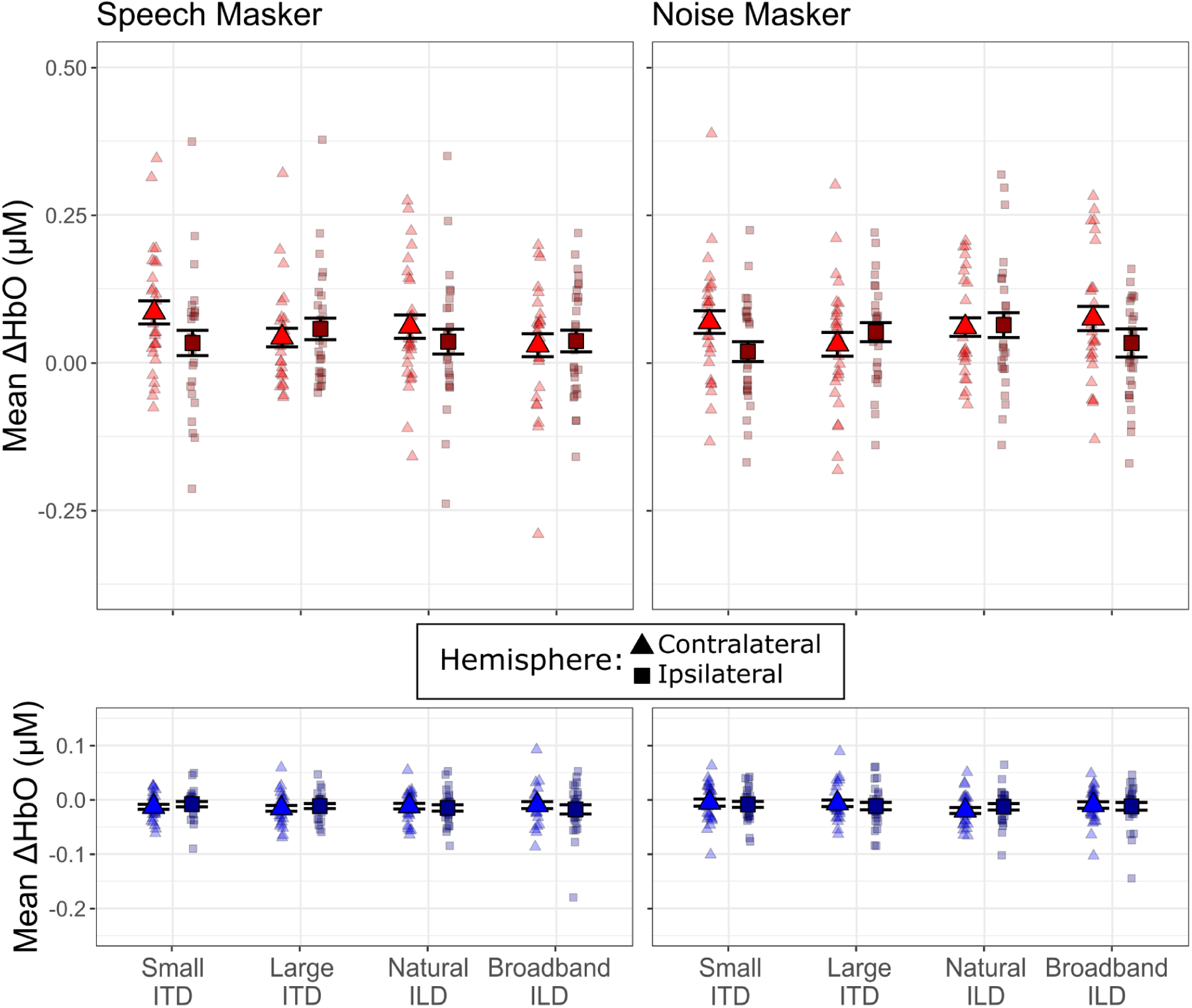
Mean hemodynamic response magnitudes in superior temporal gyrus. Top row: Mean ΔHbO when the masker was speech (left panel) and noise (right panel). Data within each spatialization condition are shown for sensors contralateral (lighter triangles) and ipsilateral (darker squares) to the direction of the target sound stream. Bottom row: As in the top row, but for mean ΔHbR. Small markers show individual participant data, and large markers show the mean across participants. Error bars indicate the mean and SEM across subjects in each condition.

These observations were supported by our statistical analyses. The mean ΔHbO models showed a significant interaction between spatialization and hemisphere both when the masker was speech (χ^2^(3) = 7.99, p = 0.046) and when the masker was noise (χ^2^(3) = 11.02, p = 0.012). To follow up on this interaction, we conducted post hoc tests comparing across hemispheres within each spatialization condition. For the speech masker, the interaction effect was driven by an effect of hemisphere in the small ITD condition only, with larger mean ΔHbO in contralateral channels (mean: 0.078 μM) compared to the ipsilateral channels (mean: 0.045 μM; β = −0.033, p = 0.0417). A similar pattern emerged in the noise masker data, with larger responses in contralateral channels (mean: 0.0645 μM) than in ipsilateral channels (mean: 0.0285 μM; β = −0.036, p = 0.017) for the small ITD condition only; the effect of hemisphere approached significance in the broadband ILD condition (contralateral mean: 0.0673, ipsilateral mean: 0.0406; β = −0.027, p = 0.077).

## Discussion

We examined how ITDs and ILDs shaped both perceptual performance and cortical activity in a color word detection task; to force listeners to rely on spatial cues, the two streams were spoken by the same talker and the timing of stimuli within each stream was irregular. Accordingly, participants were forced to segregate and select based on spatial cues. Differences in spatial cues across frequency interact with this process (Best et al., 2007), leading to patterns of behavior and neural activation.

Performance was generally better for a noise masker than a speech masker, for leading color words than lagging color words, and for stronger spatial cues (large ITDs, broadband ILDs) versus less robust spatial differences (small ITDs and natural ILDs). When listening with a speech distractor with small ITDs, hemodynamic activity in PFC was stronger than for other conditions. For both speech and noise maskers, activity in STG was lateralized contralateral to the direction of attention when sources were lateralized with small ITDs, but not for any other spatial conditions. Together, these results provide new insights into how different spatial cues contribute to SRM and speech understanding in noisy settings.

### Energetic masking accounts for noise masker results

In all spatial conditions, the hit rates were higher on average when the masker consisted of white noise bursts rather than speech. Hit rates were near ceiling, indicating that the task was relatively easy; however, rates still varied with masker type and spatialization. This supports the idea that participants were better able to hear out the target words when the background was spectrotemporally distinct from the target speech. This effect is well documented: Listeners perform better on SRM tasks when the masker is noise than when it is speech, even at negative signal to noise ratios (Best et al., 2012; Johnstone and Litovsky, 2006; Moore, 2008; Zhang et al., 2021a). Noise maskers do not contain any structured spectrotemporal features or semantic or linguistic content, and thus are not confused with words in the target stream. With a noise masker, both target segregation and selection are relatively trivial, making spatial information redundant. It is also the case that, for these stimuli, the flat-spectrum white noise bursts led to less masking in lower frequencies than did the word token maskers of equal root-mean-square intensity. The low frequencies contained both speech information and ITD cues and, in the Broadband ILD condition, substantial ILD information. Compared to the speech masker, the noise masker thus produced less energetic masking in addition to causing less interference with target speech segregation and selection.

When segregation and selection are easy (i.e., when the masker is noise), energetic masking is likely the limiting factor on performance. In these conditions, hit rates were larger for leading color words than for lagging color words. This asymmetry likely reflects the fact that the lagging token (word or noise) onset was energetically masked by the end of the leading token in each pair. Specifically, syllable onsets convey more lexical information than syllable codas (Pimentel et al., 2021; Sun and Poeppel, 2023); therefore, energetic masking of the onset of a lagging word has a greater impact on performance than masking of the coda of a leading word. For the noise masker, hit rates were lower with small ITDs than with large ITDs, which might be explained by the influence that robust ITD differences can have in combatting energetic masking (Culling and Lavandier, 2021; Durlach, 1963).

For the speech masker, hit rates were higher for leading than for lagging color words. This difference mirrors the effect for noise maskers, suggesting that energetic masking of the onsets of lagging target words degrades performance for speech maskers like it does for noise maskers. However, for speech maskers, hit rates were generally lower than for the noise masker and also varied with spatial cues, hinting that other factors affect performance, as well. These other factors are considered in the following sections.

### ITDs and ILDs yield different modes of failure for speech maskers

We hypothesized that both the small ITD and natural ILD conditions would produce relatively weak spatial cues, which could not only make it difficult to segregate target from masker but also to differentiate target words from masker words. In the small ITD condition, listeners had relatively low hit rates but also low false alarm rates: Listeners struggled to identify color words in either stream, suggesting that they struggled to group spectrotemporal features into intelligible words (i.e., a failure of segregation). This is schematized in the first row of Fig. 8.

**Figure 8.**
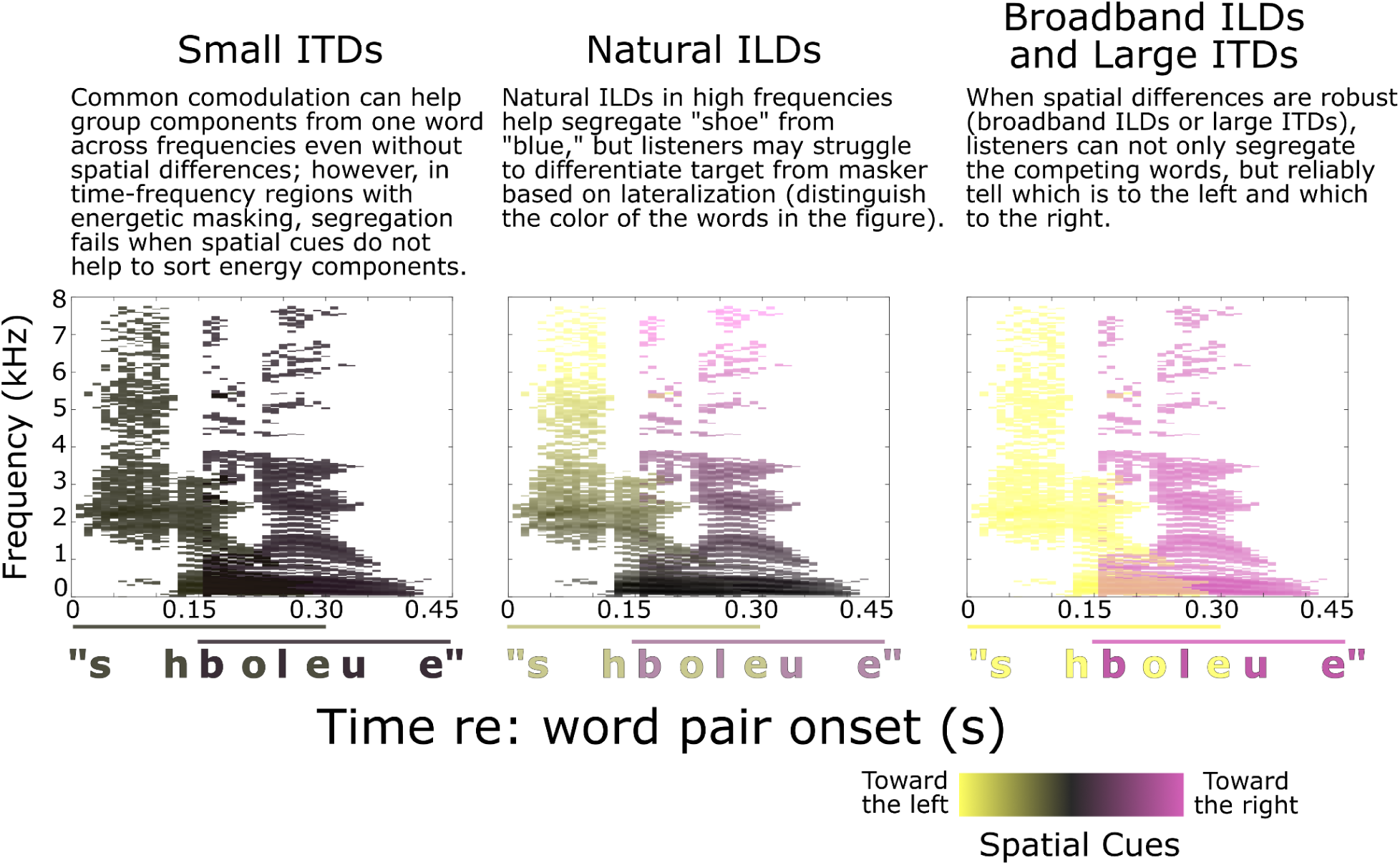
Schematic representation of how different spatial cues impact segregation and spatial perception. Each panel shows a spectrogram representation of the word pair “shoe” (leading) and “blue” (lagging). In each time-frequency pixel, the dominant spatial cues determine the color (yellow towards the left, magenta to the right). Time-frequency pixels where there is significant overlap between words are colored by the mean of the two color schemes. The horizontal lines at the bottom of each panel denote the time extent of the lead and lag words; the color of the lines and the corresponding word labels denote the perceived laterality of the overall word, integrating spatial information across the grouped spectrum. Small ITDs may fail to support segregation, leading listeners to hear an unintelligible mixture of both words (orthographically, “shboleue,” left panel). Natural ILDs may support segregation, but the perceived words may be perceived near midline, making it hard to select the target and ignore the masker. Robust spatial cues (broadband ILDs and large ITDs) enable both segregation and spatial selection.

Without access to strong spatial cues (space is conveyed with color in Fig. 8), a listener would struggle to separate “shoe” from “blue;” instead, they may have sometimes heard a mixture of both words that was difficult to interpret (“shboleue”). This also helps explain why the small ITD condition yielded a larger hemodynamic response in PFC than the other spatialization conditions; listeners expended greater cognitive effort in their attempt to segregate the competing words (Lawrence et al., 2018; Zhang et al., 2021b; Zhou et al., 2022a, 2022b). With small ITDs, activity in STG also lateralized, increasing in the hemisphere contralateral to the direction of attention — as is often observed when spatial auditory attention is strongly deployed (Alho et al., 1998, 1999, 2003; Ciaramitaro et al., 2007; Jäncke and Shah, 2002; Yang and Mayer, 2014).

In contrast, natural ILDs elicited both high hit rates and high false alarm rates. This pattern suggests that listeners could hear out individual word tokens, but could not differentiate whether color words belonged to the target or masker stream — a failure of selection, rather than segregation. Unlike in the small ITD case, with natural ILDs the two streams were discriminable from one another and listeners could hear out individual color words. However, they failed to identify whether words were from the target or the masker stream. As illustrated in the middle panel of Figure 8, ILDs in the high frequencies may help support segregation by binding with comodulated lower frequency sound components; however, the perceived lateralities of the resulting words are dominated by the lower-frequency energy, which carried zero-μs ITDs and only modest ILDs in the natural ILD condition. This is consistent with previous investigations: A low frequency interferer that is perceptually grouped with a high frequency target dominates the perceived lateral location of the composite perceptual object, (Best et al., 2007). Here, all of the competing words were perceived near midline due to the dominant zero-μs ITDs, making it hard to discriminate target from masker.

Both target and masker words are thus likely to be heard near midline, leading to increased confusions about which word was from the target and which from the masker. Notably, the false alarm rate is higher for lagging than leading masker color words, suggesting that the lateralization of the lagging word is especially weak. A number of studies have shown that interaural differences at sound onsets strongly influence lateralization (Freyman and Zurek, 2017; Klein-Hennig et al., 2011). Given that the coda of the leading word energetically masks the onset of the lagging word, it is unsurprising that listeners may be more unsure of the lateralization of a lagging color masker word than a leading color masker word, leading to an even larger false alarm rate.

Although performance was relatively poor with natural ILDs, PFC activity was not stronger in this condition than with other spatializations. In the conditions in which segregation was easier (natural ILD, large ITD, and broadband ILD), PFC was less strongly engaged and STG activity less lateralized. Together, these results hint that PFC is more strongly engaged when listeners are struggling to segregate sources, but not when they are having difficulty discriminating differences in competing stream locations (for spatial selection). For small ITDs, strong engagement of spatial attention (seen in PFC) likely increases sensory responses in contralateral STG to enhance the target stream representation, albeit with limited success. Broadband ILDs and large ITDs supported both stream segregation and spatial selection: Hit rates were high and false alarm rates were low. With large perceived spatial separation (distinct yellow and magenta in row 3 of Fig. 8), listeners both properly segregate constituent words and select the target.

These results highlight the importance of separately considering failures of segregation and spatial selection when trying to understand the processes allowing listeners to make sense of competing sounds in a mixture. Specifically, listeners must group sounds into objects based on spectrotemporal information (comodulation of different frequency components, harmonic structure, etc.). Natural ILDs alone can help support object formation of overlapping sounds by helping the brain sort out which components belong to which source. The perceived location of each object can provide a powerful cue for guiding selective attention when sources are perceived at different locations (Allen et al., 2008; Bregman, 1990; Darwin, 2007; Kidd et al., 2010; Shinn-Cunningham, 2008b). However, even spatial cues that are sufficient to aid source segregation (like natural ILDs) may not produce robust differences in perceived location, leading to confusions between properly segregated sources.

### Summary, Limitations, and Future Directions

Here, we compared behavioral performance and hemodynamic responses in ILD and ITD spatialization conditions. We found that weaker ITDs and ILDs (small and natural, respectively) led to poorer performance than stronger ITDs and ILDs (large and broadband, respectively). Moreover, differences in the response patterns suggest that small ITDs led to failures of segregation, while natural ILDs, which are modest in the frequency range where most speech energy resides, produced more selection errors. It is tempting to conclude that ITDs and ILDs thus contribute differently to SRM performance, in general. However, we did not match these “weak” ITD and ILD conditions to produce perceptually similar lateralization of our stimuli, limiting our ability to directly compare across conditions.

In the small ITD condition, we observed greater hemodynamic responses in PFC and a lateralization of hemodynamic responses in STG, with greater activation in the hemisphere contralateral to the direction of the target. Considered alongside our behavioral results, this pattern suggests that PFC was more strongly engaged in the one condition where segregation failures limited performance. Although these findings are intriguing, they are not definitive. The measures of hemodynamic activity obtained with this fNIRS montage were relatively coarse with four channels of data per hemisphere of STG and seven channels of data in PFC. We also gathered only eight blocks of data per condition. Further investigations employing denser fNIRS montages, presenting more blocks in each spatialization condition, and co-registering results with MRI structural scans would provide more definitive results (Yücel et al., 2024). Such studies would provide better ground-truth estimates of the neural locus of activity with better signal-to-noise ratio.

Future experiments should investigate psychophysically matched ITDs and ILDs or parametrically vary ITD and ILD magnitudes to support direct comparisons between ITDs and ILDs that produce similar spatial percepts. Clinically focused experiments could also explore whether improved spatial hearing outcomes in bilateral cochlear implant users with magnified ILDs (Brown, 2014, 2018; Richardson et al., 2025) are driven by the same mechanisms as in normal hearing listeners.

